## Supplementary material for "Novel MOG analogues to explore the MCT2 pharmacophore, α-ketoglutarate biology and cellular effects of *N*-oxalylglycine": Chemical synthesis methods

### Synthesis of MOG Analogues

#### 2-((2-Methoxy-2-oxoethyl)amino)-2-oxoacetic acid (MOG, 1)

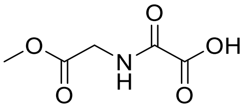

This compound was prepared according to the procedure described in Fets *et al.*, *Nat. Chem. Biol.* 2018, 14, 1032–1042.

#### *tert*-Butyl 2-chloro-2-oxoacetate (General Intermediate)

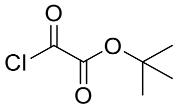

A stirred solution of oxalyl chloride (5.08 g, 40 mmol) in $Et_2O$ (80 mL) was cooled at 0 °C and *tert*-butanol (2.97 g, 40 mmol) in $Et_2O$ (20 mL) was added dropwise under $N_2$. The resulting mixture was stirred for 30 min at 0 °C and then 16 h at room temperature. The solvent was evaporated to afford *tert*-butyl 2-chloro-2-oxoacetate a colourless oil. This material was used in crude form for the preparation of each of the intermediate described below.

#### *tert*-Butyl 2-(2-ethoxy-2-oxoethylamino)-2-oxoacetate (Intermediate 2)

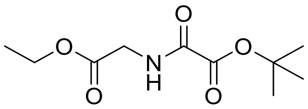

*tert*-Butyl 2-chloro-2-oxoacetate (712 mg, 4.32 mmol) was dissolved in $CH_2Cl_2$ (5 mL) and added dropwise to a solution of glycine ethyl ester hydrochloride (604 mg, 4.32 mmol) and $Et_3N$ (874 mg, 8.65 mmol) in $CH_2Cl_2$ (20 mL) under $N_2$ at 0 °C. The resulting mixture was stirred from 0 °C to room temperature for 3 h. The solvent was evaporated under reduced pressure and the residue was purified by chromatography on silica eluting with petrol ether:EtOAc (3:1) to afford the title product (738 mg, 73%) as a clear oil. **LCMS** Rt = 1.35 min, $[2M+Na]^+$ = 485.3; **$^1$H NMR**.(400 MHz, $CD_3OD$) δ 4.22 (q, $J$ = 8.0 Hz, 2H), 4.03 – 4.00 (m, 2H), 1.58 (s, 9H), 1.29 (t, $J$ = 8.0 Hz, 3H).

#### 2-((2-Ethoxy-2-oxoethyl)amino)-2-oxoacetic acid (2)

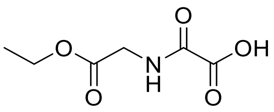

A mixture of *tert*-butyl 2-(2-ethoxy-2-oxoethylamino)-2-oxoacetate (728 mg, 3.15 mmol) and $CF_3COOH$ (3 mL) in $CH_2Cl_2$ (10 mL) was stirred at room temperature for 3 h. The solvent was evaporated under reduced pressure. $Et_2O$ was added to the residue and removed under reduced pressure to afford the title product (388 mg, 100%) as an off-white solid. **$^1$H NMR** (400 MHz, DMSO-d$_6$) δ 9.10 (t, *J* = 6.0 Hz, 1H), 4.11 (q, *J* = 7.2 Hz, 2H), 3.87 (d, *J* = 6.0 Hz, 2H), 1.19 (t, *J*=7.2 Hz, 3H).

**_tert_-Butyl 2-(2-isopropoxy-2-oxoethylamino)-2-oxoacetate (Intermediate 3)**

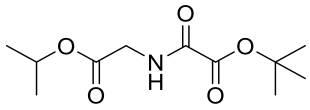

*tert*-Butyl 2-chloro-2-oxoacetate (1.074 g, 6.53 mmol) was dissolved in $CH_2Cl_2$ (10 mL) and added dropwise to a solution of glycine isopropyl ester hydrochloride (1.00 g, 6.53 mmol) and $Et_3N$ (1.32 g, 13 mmol) in $CH_2Cl_2$ (20 mL) under $N_2$ at 0 °C. The resulting mixture was stirred from 0 °C to room temperature for 3 h. The solvent was evaporated under reduced pressure and the residue was purified by chromatography on silica eluting with petrol ether:EtOAc (3:1) to afford the title product (212 mg) as a clear oil. **LCMS** Rt = 1.43 min, $[2M+Na]^+$ = 513.3; **$^1$H NMR**.(400 MHz, $CDCl_3$) δ 7.51 (s, 1H), 5.12 (spt, *J* =6.0 Hz, 1H), 4.06 (d, *J* = 6.0 Hz, 2H), 1.59 (s, 9H), 1.30 (d, *J* = 6.0 Hz, 6H).

**2-((2-Isopropoxy-2-oxoethyl)amino)-2-oxoacetic acid (3, IPOG)**

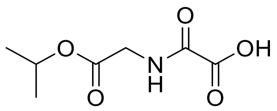

A mixture of *tert*-butyl 2-(2-isopropoxy-2-oxoethylamino)-2-oxoacetate (132 mg, 0.54 mmol) and $CF_3COOH$ (3 mL) in $CH_2Cl_2$ (10 mL) was stirred at room temperature for 3 h. The solvent was evaporated under reduced pressure and the solid freeze dried to afford the title product (52 mg, 51%) as a white solid. **$^1$H NMR** (400 MHz, DMSO-d$_6$) δ = 9.07 (t, *J* = 6.0 Hz, 1H), 4.92 (spt, *J*=6.3 Hz, 1H), 3.83 (d, *J*=6.0 Hz, 2H), 1.19 (d, *J*=6.3 Hz, 6H).

**(_R_)-Methyl 2-(2-(_tert_-butoxy)-2-oxoacetamido)propanoate (Intermediate 4)**

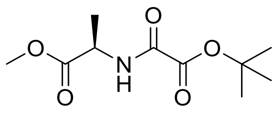

(*R*)-methyl 2-aminopropanoate hydrochloride (746 mg, 5.34 mmol) was dissolved in $CH_2Cl_2$ (5 mL) and $Et_3N$ (1.08 g, 10.7 mmol) was slowly added. The mixture was stirred at 0 °C for 10 min. *tert*-Butyl 2-chloro-2-oxoacetate (800 mg, 4.86 mmol) was added dropwise and the resulting mixture was stirred from 0 °C to room temperature for 3 h. The solvent was evaporated under reduced pressure and the residue was purified by chromatography on silica eluting with petrol ether:EtOAc (3:1) to afford the title product (520 mg, 46%) as a white solid.

**[1]H NMR** (400 MHz, $CD_3OD$) δ 4.49 (q, *J* = 8.0 Hz, 1H), 3.76 (s, 3H), 1.58 (s, 9H), 1.46 (d, *J* = 8.0 Hz, 3H).

**(*R*)-Methyl 2-aminopropanoate (4)**

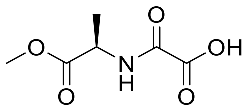

A mixture of (*R*)-methyl 2-(2-(*tert*-butoxy)-2-oxoacetamido)propanoate (740 mg, 2.24 mmol) and $CF_3COOH$ (3 mL) in $CH_2Cl_2$ (10 mL) was stirred at room temperature for 3 h. The solvent was evaporated under reduced pressure and the solid was dried by lyophilization to afford the title product (560 mg, 100%) as an off-white solid. **[1]H NMR** (400 MHz, DMSO-d6) δ 9.12 (d, *J* = 7.3 Hz, 1H), 4.34 (quin, *J* = 7.4 Hz, 1H), 3.63 (s, 3H), 1.34 (d, *J* = 7.3 Hz, 3H).

**(*S*)-Methyl 2-(2-(*tert*-butoxy)-2-oxoacetamido)propanoate (Intermediate 5)**

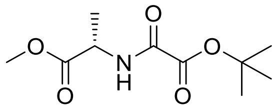

*tert*-Butyl 2-chloro-2-oxoacetate (368 mg, 2 mmol) in $CH_2Cl_2$ (2 mL) was added dropwise to a solution of (*S*)-methyl 2-aminopropanoate hydrochloride (284 mg, 2 mmol) and $Et_3N$ (267 mg, 2.24 mmol) in $CH_2Cl_2$ (5 mL) at 0 °C. The resulting mixture was stirred from 0 °C to room temperature for 3 h. The solvent was evaporated under reduced pressure and the residue was purified by chromatography on silica eluting with petrol ether:EtOAc (4:1) to afford the title product (112 mg, 24%) as a colourless oil. **[1]H NMR** (400 MHz, $CDCl_3$) δ 7.59 (s, 1H), 4.63 – 4.59 (m, 1H), 3.80 (s, 3H), 1.59 (s, 9H), 1.49 (d, *J* = 8.0 Hz, 3H).

**(*S*)-Methyl 2-aminopropanoate (5)**

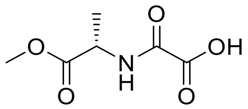

A mixture of (*S*)-methyl 2-(2-(*tert*-butoxy)-2-oxoacetamido)propanoate (112 mg, 0.48 mmol) and $CF_3COOH$ (3 mL) in $CH_2Cl_2$ (10 mL) was stirred at room temperature for 3 h. The solvent was evaporated under reduced pressure to afford the title product (83 mg, 99%) as a white solid. **[1]H NMR** (400 MHz, DMSO-$d_6$) δ 9.09 (d, *J* = 7.3 Hz, 1H), 4.34 (quin, *J* = 7.3 Hz, 1H), 3.64 (s, 3H), 1.34 (d, *J* = 7.3 Hz, 3H).

**Methyl 2-(2-(*tert*-butoxy)-2-oxoacetamido)-2-methylpropanoate (Intermediate 6)**

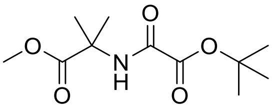

*tert*-Butyl 2-chloro-2-oxoacetate (591 mg, 3.6 mmol) in $CH_2Cl_2$ (10 mL) was added to a solution of methyl 2-amino-2-methylpropanoate (500 mg, 3.24 mmol) and $Et_3N$ (690 mg, 6.83 mmol) in $CH_2Cl_2$ (20 mL) at 0 °C under a nitrogen atmosphere. The resulting mixture was stirred for 5 min at 0 °C and at room temperature for 2 h. Saturated aqueous $NaHCO_3$ (30 mL) was added, the organic phase was separated, dried over $Na_2SO_4$, filtered and evaporated under reduced pressure to afford the crude title product as a yellow oil (870 mg, >100%). This material was taken into deprotection step without purification.

**2-((1-Methoxy-2-methyl-1-oxopropan-2-yl)amino)-2-oxoacetic acid (6)**

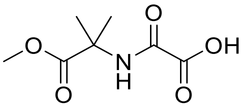

To a stirred solution of methyl 2-(2-(*tert*-butoxy)-2-oxoacetamido)-2-methylpropanoate (870 mg, 3.55 mmol) in $CH_2Cl_2$ (21.3 mL) was added $CF_3COOH$ (7 mL). The reaction mixture was stirred at room temperature for 7 h. The solvent was evaporated under reduced pressure. The residue was purified by HPLC (Xbridge Prep C18 OBD column 19 x 150 mm 5 µm; Mobile Phase A: Water (0.1% $CF_3COOH$), Mobile Phase B: acetonitrile; flowrate 20 mL/min; gradiant 5% B to 25.6% B in 5 min; UV detection 254 nm) to afford the title product (257 mg, 24%) as a white solid after freeze drying. **LCMS** Rt = 1.07 min, $[MH]^+$ = 190.0; $^1$**H NMR** (400 MHz, DMSO-$d_6$) δ 13.97 (s, 1H), 8.99 (s, 1H), 3.59 (s, 3H), 1.40 (s, 6H).

***tert*-Butyl 2-oxo-2-((2-oxopropyl)amino)acetate (Intermediate 7)**

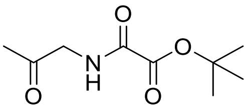

1-Aminopropan-2-one hydrochloride (431 mg, 3.93 mmol) in $CH_2Cl_2$ (20 mL) was cooled to 0 °C and $Et_3N$ (795 mg, 7.86 mmol) was added. The reaction mixture was stirred at room temperature for 10 min and then cooled to 0 °C. *tert*-Butyl 2-chloro-2-oxoacetate (647 mg, 3.93 mmol) was added dropwise and the resulting mixture was stirred from 0 °C to room temperature for 3 h. The solvent was evaporated under reduced pressure and the residue was purified by chromatography on silica eluting with petrol ether:EtOAc (1:1) to afford the title product (71 mg, 9%) as a yellow oil. **LCMS** Rt = 1.22 min, $[2M+Na]^+$ = 425.3.

**2-Oxo-2-((2-oxopropyl)amino)acetic acid (7)**

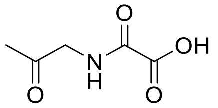

A mixture of *tert*-butyl 2-oxo-2-((2-oxopropyl)amino)acetate (71 mg, 0.35 mmol) and $CF_3COOH$ (2 mL) in $CH_2Cl_2$ (6 mL) was stirred at room temperature for 3 h. The solvent was evaporated under reduced pressure to afford the title product (51 mg, 100%) as an off-white solid. **$^1$H NMR** (400 MHz, DMSO-$d_6$) δ 8.84 (br. s., 1H), 3.98 (d, *J* = 6.0 Hz, 2H), 2.09 (s, 3H).

***tert*-Butyl 2-((oxazol-2-ylmethyl)amino)-2-oxoacetate (Intermediate 8)**

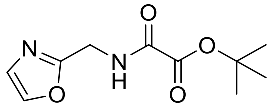

Oxazol-2-ylmethanamine hydrochloride (204 mg, 2 mmol) and $Et_3N$ (420 mg, 4.16 mmol) in $CH_2Cl_2$ (20 mL) were cooled to 0 °C and stirred for 20 min. *tert*-Butyl 2-chloro-2-oxoacetate (342 mg, 2 mmol) in $CH_2Cl_2$ (10 mL) was added dropwise and the resulting mixture was stirred from 0 °C to room temperature for 3 h. The solvent was evaporated under reduced pressure and the residue was purified by chromatography on silica eluting with petrol ether:EtOAc (1:1) to afford the title product (187 mg, 71%) as a white solid. **$^1$H NMR** (400 MHz, $CDCl_3$) δ 7.67 (s, 2H), 7.12 (s, 1H), 4.67 (d, *J* = 6.0 Hz, 2H).

**2-((Oxazol-2-ylmethyl)amino)-2-oxoacetic acid (8)**

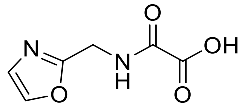

A mixture of *tert*-butyl 2-((oxazol-2-ylmethyl)amino)-2-oxoacetate (187 mg, 0.83 mmol) and $CF_3COOH$ (3 mL) in $CH_2Cl_2$ (10 mL) was stirred at room temperature for 3 h. The solvent was evaporated under reduced pressure to afford the title product (135 mg, 96%) as a white solid. **$^1$H NMR** (400 MHz, DMSO-$d_6$) δ 9.35 (t, *J* = 6.0 Hz, 1H), 8.03 (s, 1H), 7.15 (s, 1H), 4.43 (d, *J* = 6.0 Hz, 2H).

***tert*-Butyl 2-oxo-2-(((5-(trifluoromethyl)furan-2-yl)methyl)amino)acetate (Intermediate 9)**

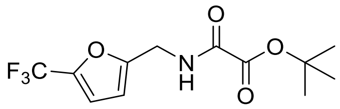

*tert*-Butyl 2-chloro-2-oxoacetate (134 mg, 0.82 mmol) in $CH_2Cl_2$ (5 mL) was added to a solution of (5-(trifluoromethyl)furan-2-yl)methanamine hydrochloride (150 mg, 0.746 mmol) and $Et_3N$ (158 mg, 1.574 mmol) in $CH_2Cl_2$ (25 mL) at 0 °C under nitrogen atmosphere. The resulting mixture was stirred for 5 min at 0 °C and at room temperature for 2 h. Saturated aqueous

$NaHCO_3$ (30 mL) was added, the organic phase was separated, dried over $Na_2SO_4$, filtered and evaporated under reduced pressure to afford the crude title product as a white solid (567 mg, >100%). This material was taken into deprotection step without purification.

**2-Oxo-2-(((5-(trifluoromethyl)furan-2-yl)methyl)amino)acetic acid (9)**

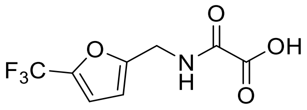

To a stirred solution of *tert*-Butyl 2-oxo-2-(((5-(trifluoromethyl)furan-2-yl)methyl)amino)acetate (567 mg, 1.935 mmol) in $CH_2Cl_2$ (12 mL) was added $CF_3COOH$ (3.87 mL). The reaction mixture was stirred at room temperature for 7 h. The solvent was evaporated under reduced pressure. The residue was purified by HPLC HPLC (Xbridge Prep C18 OBD column 19 x 150 mm 5 µm; Mobile Phase A: Water (0.1% $CF_3COOH$), Mobile Phase B: acetonitrile; flowrate 20 mL/min; gradiant 1% B to 100% B in 3 min; UV detection 220 nm) to afford the title product (73 mg, 16%) as a white solid after freeze drying. **LCMS** Rt = 0.66 min, $[M-H]^-$ = 236.0; **$^1$H NMR** (300 MHz, DMSO-$d_6$) δ 13.99 (s, 1H), 9.39 (t, *J* = 5.4 Hz, 1H), 7.15-7.14 (m, 1H), 6.49-6.48 (m, 1H), 4.38 (d, *J* = 6 Hz, 2H).

***tert*-Butyl 2-(((3-methyl-1,2,4-oxadiazol-5-yl)methyl)amino)-2-oxoacetate (Intermediate 10)**

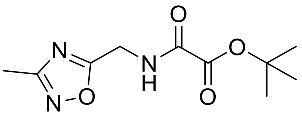

*tert*-Butyl 2-chloro-2-oxoacetate (182 mg, 1.11 mmol) in $CH_2Cl_2$ (5 mL) was added to a solution of (3-methyl-1,2,4-oxadiazol-5-yl)methanamine hydrochloride (150 mg, 1 mmol) and $Et_3N$ (213 mg, 2.11 mmol) in $CH_2Cl_2$ (25 mL) at 0 °C under nitrogen atmosphere. The resulting mixture was stirred for 5 min at 0 °C and at room temperature for 2 h. Saturated aqueous $NaHCO_3$ (30 mL) was added, the organic phase was separated, dried over $Na_2SO_4$, filtered and evaporated under reduced pressure to afford the crude title product as a yellow oil (238 mg, 99%). This material was taken into deprotection step without purification.

**2-(((3-Methyl-1,2,4-oxadiazol-5-yl)methyl)amino)-2-oxoacetic acid (10)**

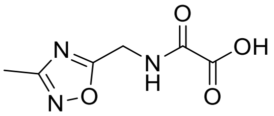

To a stirred solution of *tert*-butyl 2-(((3-methyl-1,2,4-oxadiazol-5-yl)methyl)amino)-2-oxoacetate (238 mg, 0.987 mmol) in $CH_2Cl_2$ (6 mL) at 0 °C under nitrogen atmosphere was added $CF_3COOH$ (2 mL). The reaction mixture was stirred at 0 °C for 5 min and at room temperature for 6 h. The solvent was evaporated under reduced pressure. The residue was purified by HPLC (XSelect Prep C18 OBD column 19 x 150 mm 5 µm; Mobile Phase A: Water (0.05% $CF_3COOH$), Mobile Phase B: acetonitrile; flowrate 20 mL/min; gradiant 2% B to 15% B in 7 min; UV detection 254 nm) to afford the title product (31 mg, 17%) as a white solid after freeze drying. **LCMS** Rt = 0.28 min, $[MH]^+$ = 185.9; $^1$**H NMR** (400 MHz, DMSO-$d_6$) δ 14.16 (s, 1H), 9.55 (t, *J* = 5.6 Hz, 1H), 4.57 (d, *J* = 6 Hz, 2H), 2.32 (s, 3H).
